## Supplementary information for "Neuroligin-mediated neurodevelopmental defects are induced by mitochondrial dysfunction and prevented by lutein in *C. elegans*"

#### Table of Content

1. Supplementary Figures: 1 - 12
2. Supplementary Tables: 1 - 10
3. Supplementary Methods
4. Supplementary information references

#### Supplementary Figures

##### Supplementary Figure 1

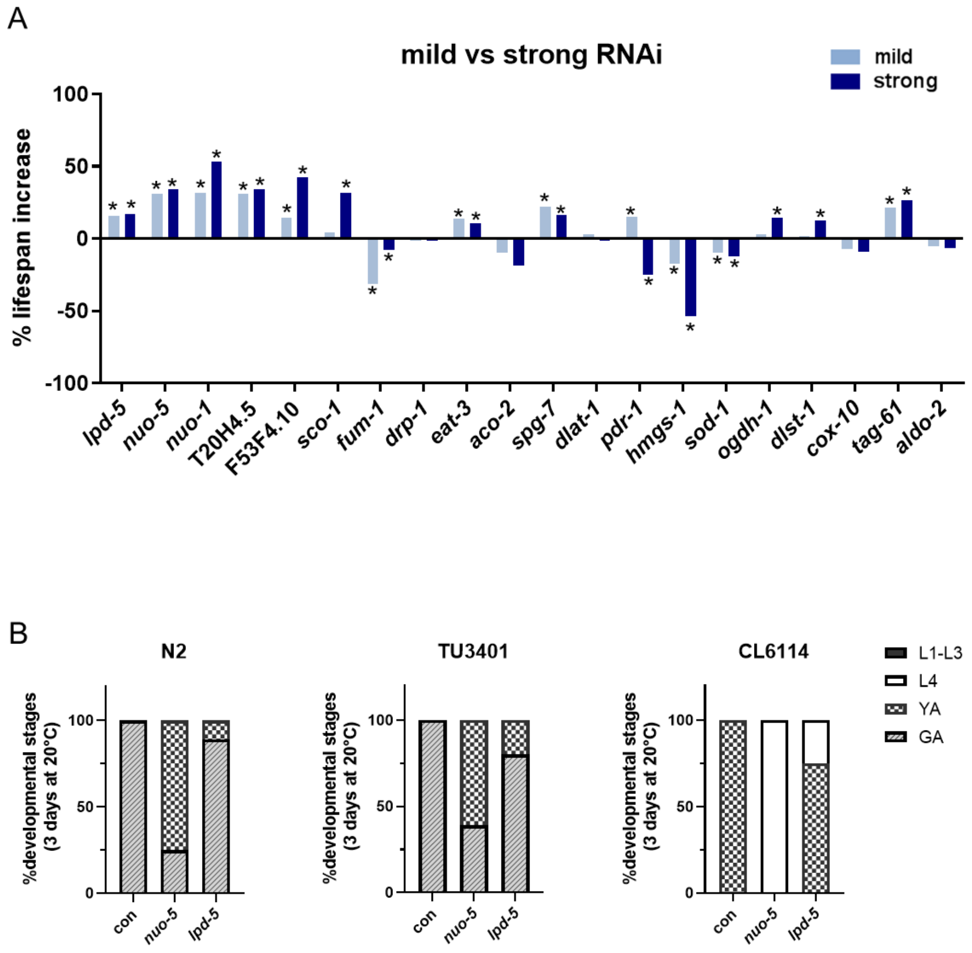


**Figure S1. Effect of RNAi against the mitochondrial proteins.**
**A)** Summary of survival analysis: % changes in mean lifespan. Wild-type (N2) worms fed bacteria expressing either *lpd-5*, *nuo-5*, T20H4.5, F53F4.10, *nuo-1*, C01F1.2, *fum-1*, *drp-1*, *eat-3*, *aco-2*, *spg-7*, *pdr-1*, *sod-1*, F23B12.5, W02F12.5, *tag-61*, F01F1.12 or F25B4.6 dsRNA. See Table S2 and S3 for summary statistics from Kaplan-Meier survival analysis. Asterisks denote statistical significance compared to control. *p-value <0.0001. Bar graphs represent % changes in means lifespan compared to N2 fed empty-vector (control)**. B)** Comparison of development between N2 (wild-type) and RNAi sensitive *C. elegans* strain. Bars indicate the percentage of different developmental stages after 3 days from egg lay. L1 -L3, 1$^{st}$ to 3$^{rd}$ larval stage; L4, 4$^{th}$ larval stage; YA, young adult; GA, gravid adult. N=2, n=40

**Supplementary Figure 2**

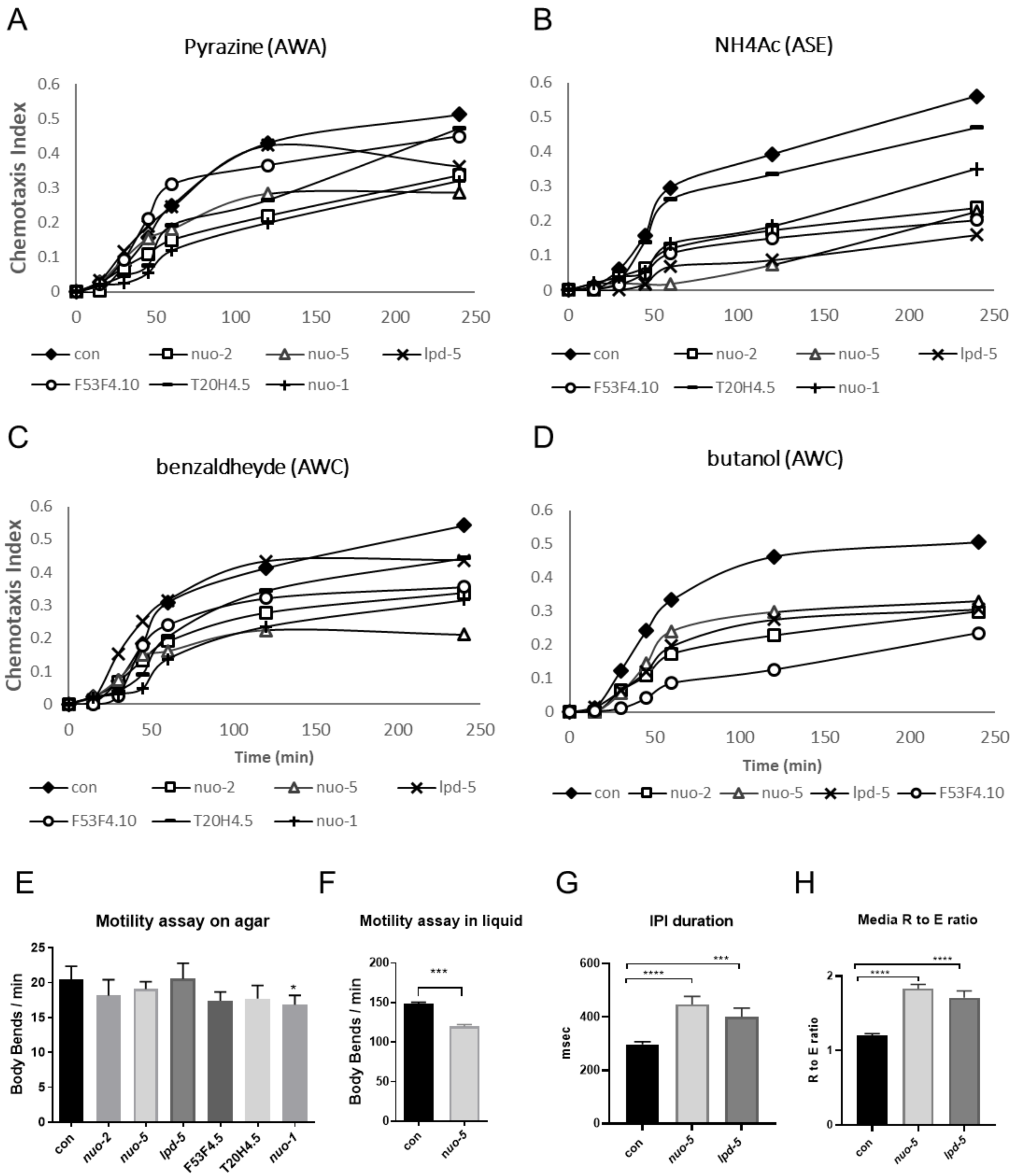


**Figure S2. Strong RNAi against the mitochondrial Complex I proteins affects chemotaxis, motility and pharyngeal pumping. A-D)** impairment in sensory perception upon strong RNAi against Complex I proteins. Chemotaxis curves obtained using 1µl of either 1% Pyrazine **(A),** 2.5M Sodium Acetate **(B),** 1% Benzaldehyde **(C)** or 1%Butanol **(D)** as attractant for the worms (two-way ANOVA followed by Tukey's multiple-comparisons test).

**E**) Motility assay on agar (one-way ANOVA) and **F)** in liquid media (Student's t-test). Locomotion is quantified as body bend per minute. (N=3, n=10-15 per group). **G**) IPI (interpump interval) duration and **H**) Median R to E ratio were measured using the ScreenChipTM System (InVivo Biosystems). Bars represent means ± SEM.

**Supplementary Figure 3**

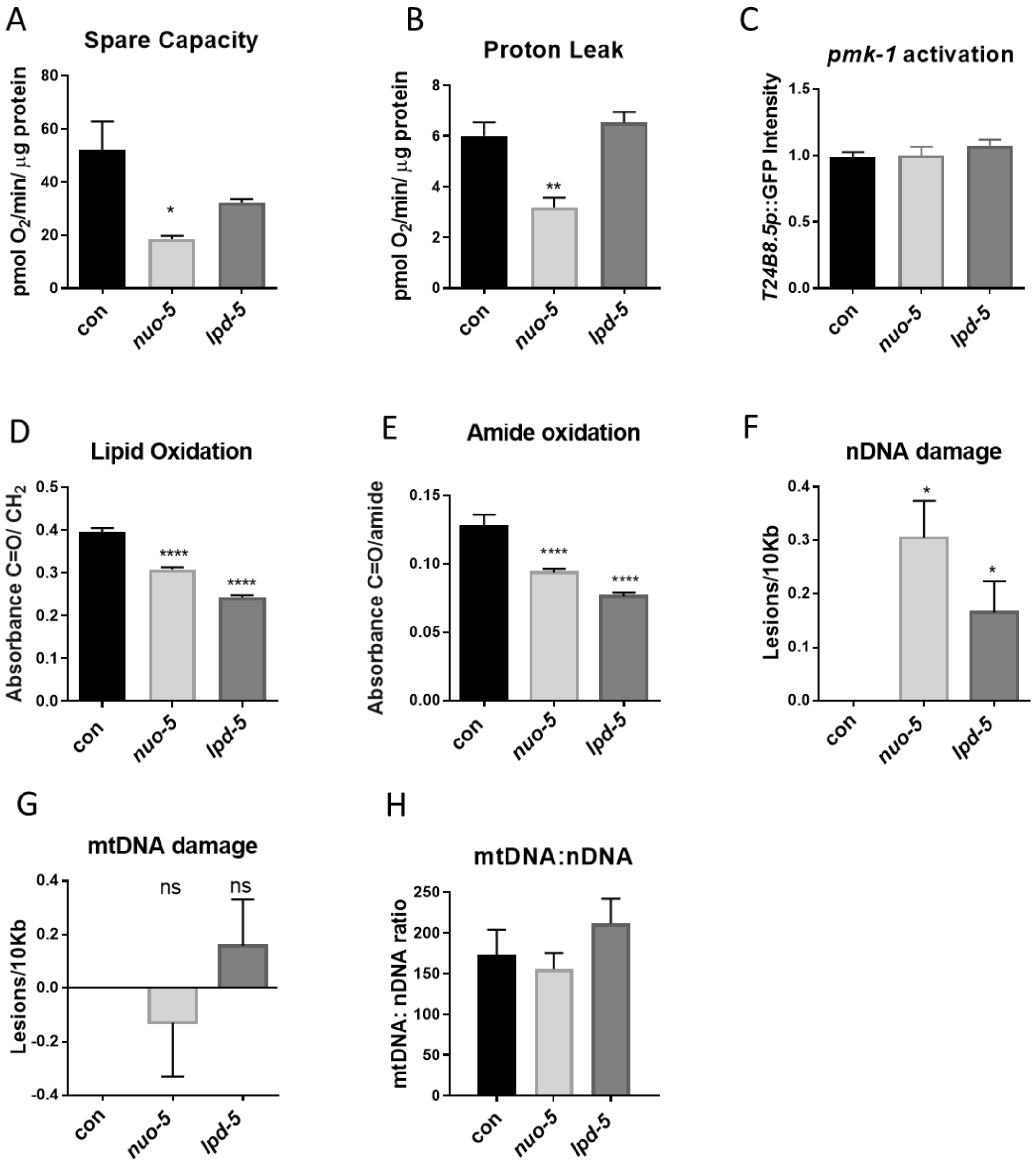


**Figure S3. Strong RNAi against the mitochondrial Complex I proteins does not lead to significant signs of oxidative damage. A**) Proton leak and **B**) Spare capacity were measured using the Seahorse XF24 Analyzer (N=3 biologically independent experiments n=1000-1500 per group). **C)** Quantification of GFP expression in a WT strains expressing the *agIs219* transgene, which is comprised of the promoter of a PMK-1-regulated gene, *T24B8.5* fused to GFP and provides an *in vivo* sensor of PMK-1 pathway activity (N=2, n=20-25). **D**) Lipid oxidation and **E**) amide oxidation assessed with SR-μFTIR. (N=3, n=15-20 per group). **F)** Nuclear DNA (nDNA) damage, **G)** Mitochondrial DNA (mtDNA) damage and **H)** mtDNA copy number upon strong suppression of *lpd-5* and *nuo-5*. In every panel worms were fed as in Fig 2 and statistical analysis was carried out using one-way ANOVA.

**Supplementary Figure 4**

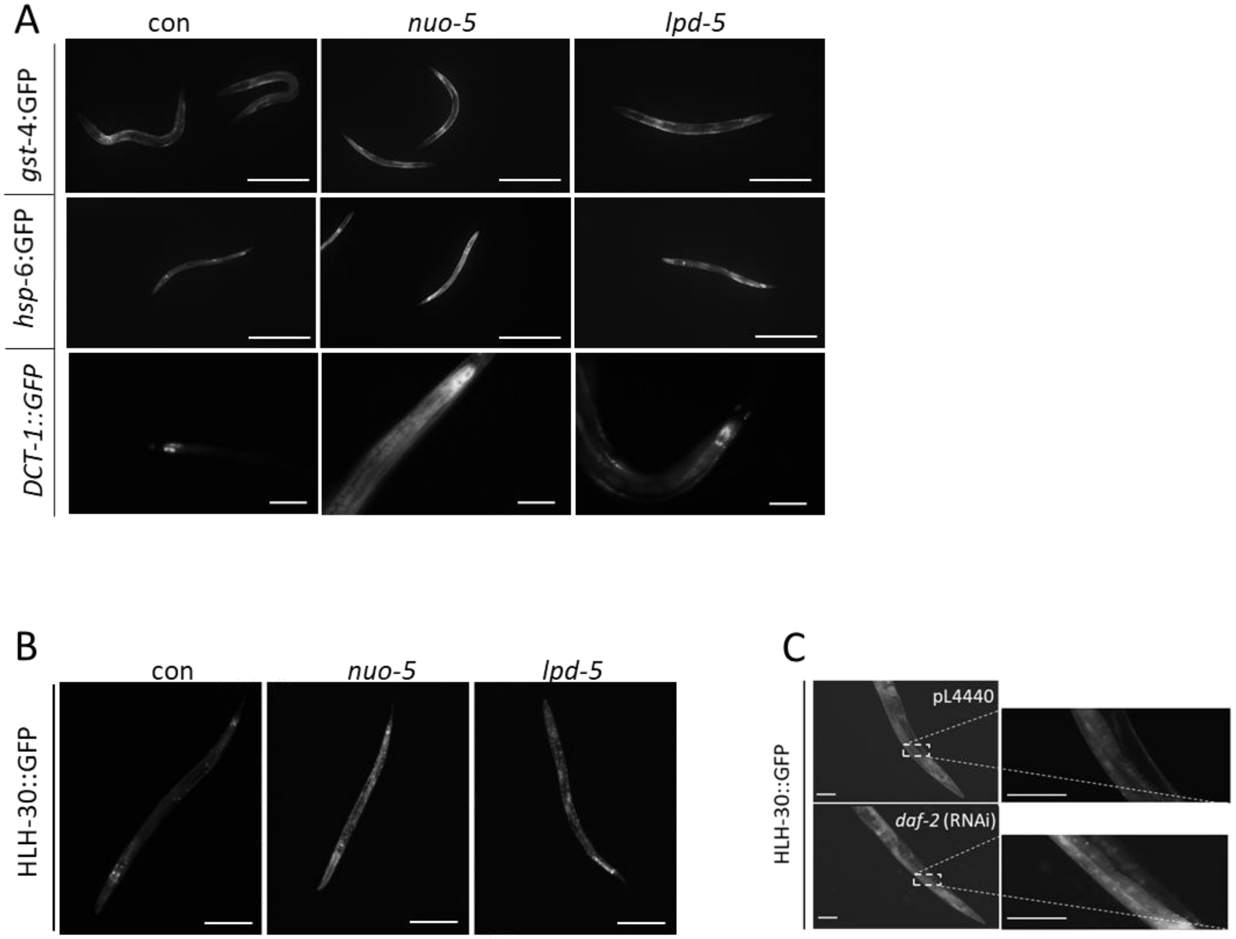


**Figure S4. Strong RNAi against the mitochondrial Complex I proteins differentially affect the expression of mitochondrial stress response genes. A**) Representative pictures of GFP reporter strains upon *nuo-5* and *lpd-5* RNAi. Scale bar 370 µm for p$_{gst\text{-}4}$::GFP and p$_{hsp\text{-}6}$::GFP and 50 µm for p$_{dct\text{-}1}$DCT-*1*::GFP. Worms are L3 larvae. **B)** Representative pictures of nuclear translocation of P$_{hlh\text{-}30}$::HLH-30::GFP upon *nuo-5 and lpd-5* RNAi. Worms are L3 larvae. Scale bars 100 µm. **C**) Representative pictures of nuclear translocation of P$_{hlh\text{-}30}$::HLH-30::GFP upon *daf-2* RNAi in one day old worms. Scale bars 100 µm.

**Supplementary Figure 5**

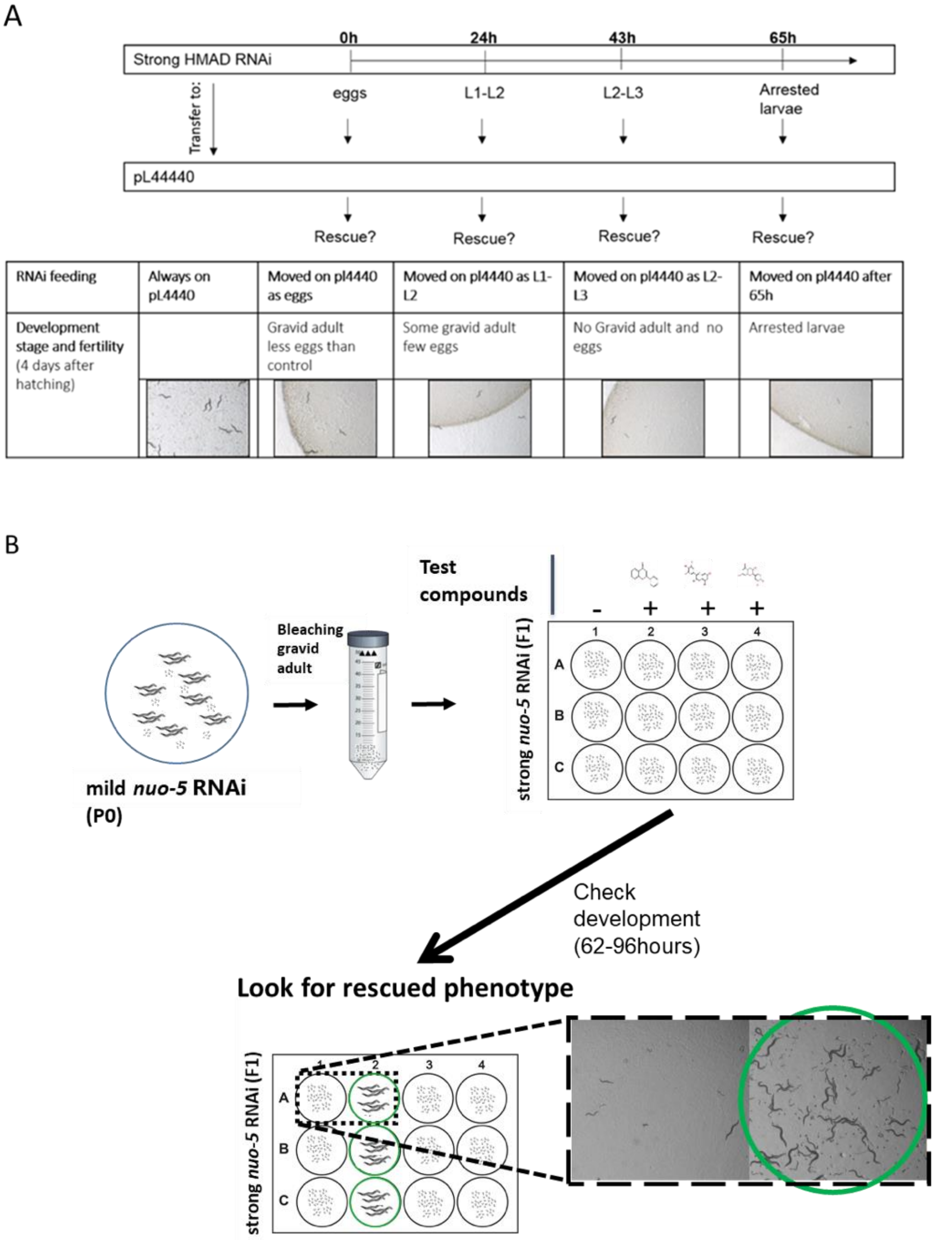


**Figure S5. Drug suppressor screening proof of principle and workflow. A**) Schematic experimental strategy to proof the reversibility of the strong phenotype. Animals were cultured on RNAi plates (strong treatment); then worms were moved on control plates (fed the empty vector pL4440) either as eggs or at different larval stages. The development was checked for rescue on day 4 after egg hatching (representative pictures of a partial rescue obtained returning the nematodes on pL4440 within 24 hours). **B**) Flowchart with representative picture of the compounds screen in search of suppressors of the *nuo-5* and *lpd-5* strong (developmental arrest) phenotype.

#### Supplementary Figure 6

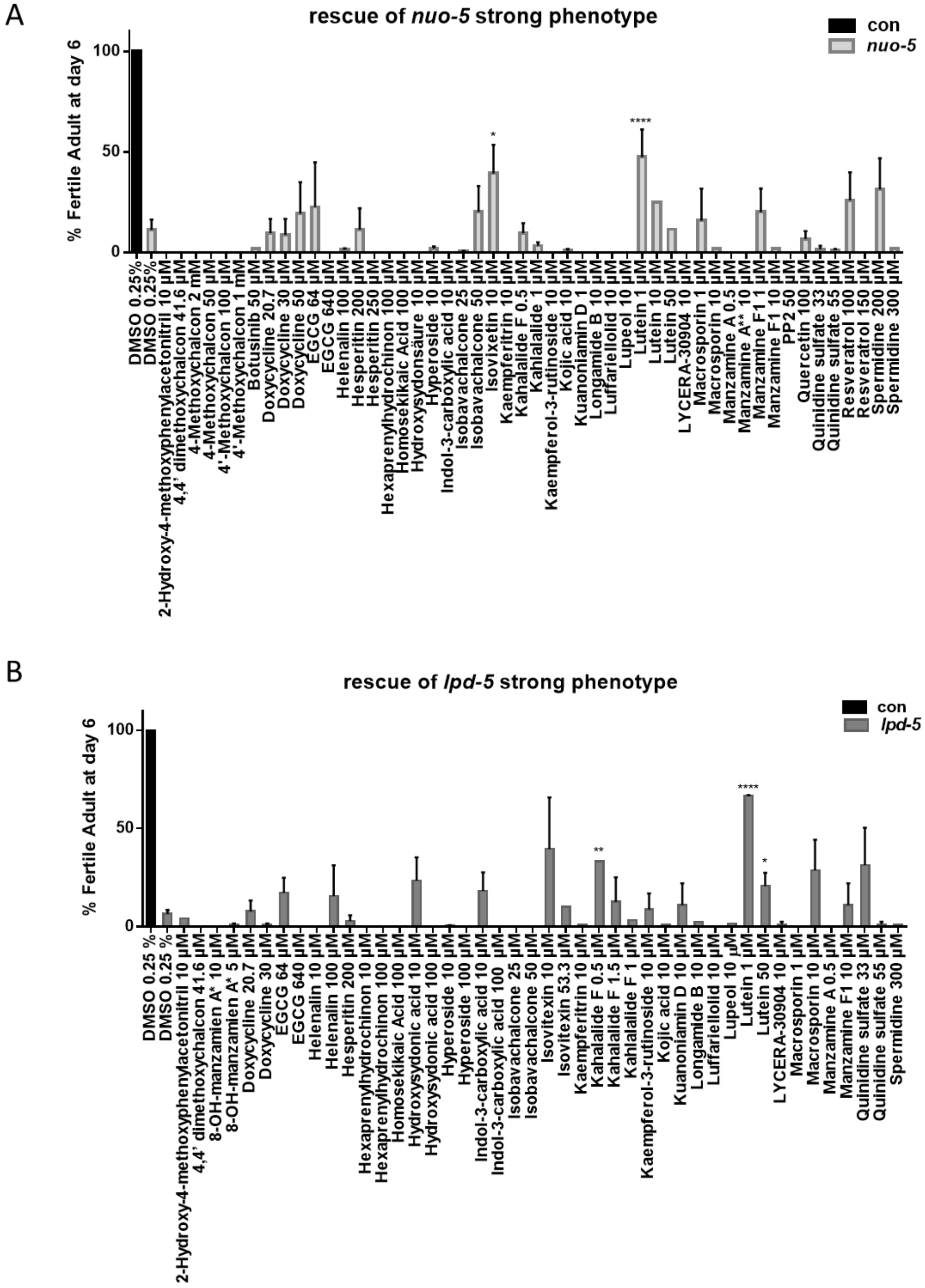


**Figure S6. Results of suppressor screening.** Bars with numbers show the percentage of animals which developed to gravid adult after 6 days from eggs ± SEM. **A)** * p-value 0.0379 and ****p-value < 0.0001 compared to *nuo-5* RNAi treated with DMSO; **B**) *p-value 0.022, ** p-value 0.001, ****p-value < 0.0001 compared to *lpd-5* RNAi treated with DMSO (n= 40, one-way ANOVA).

### Supplementary Figure 7

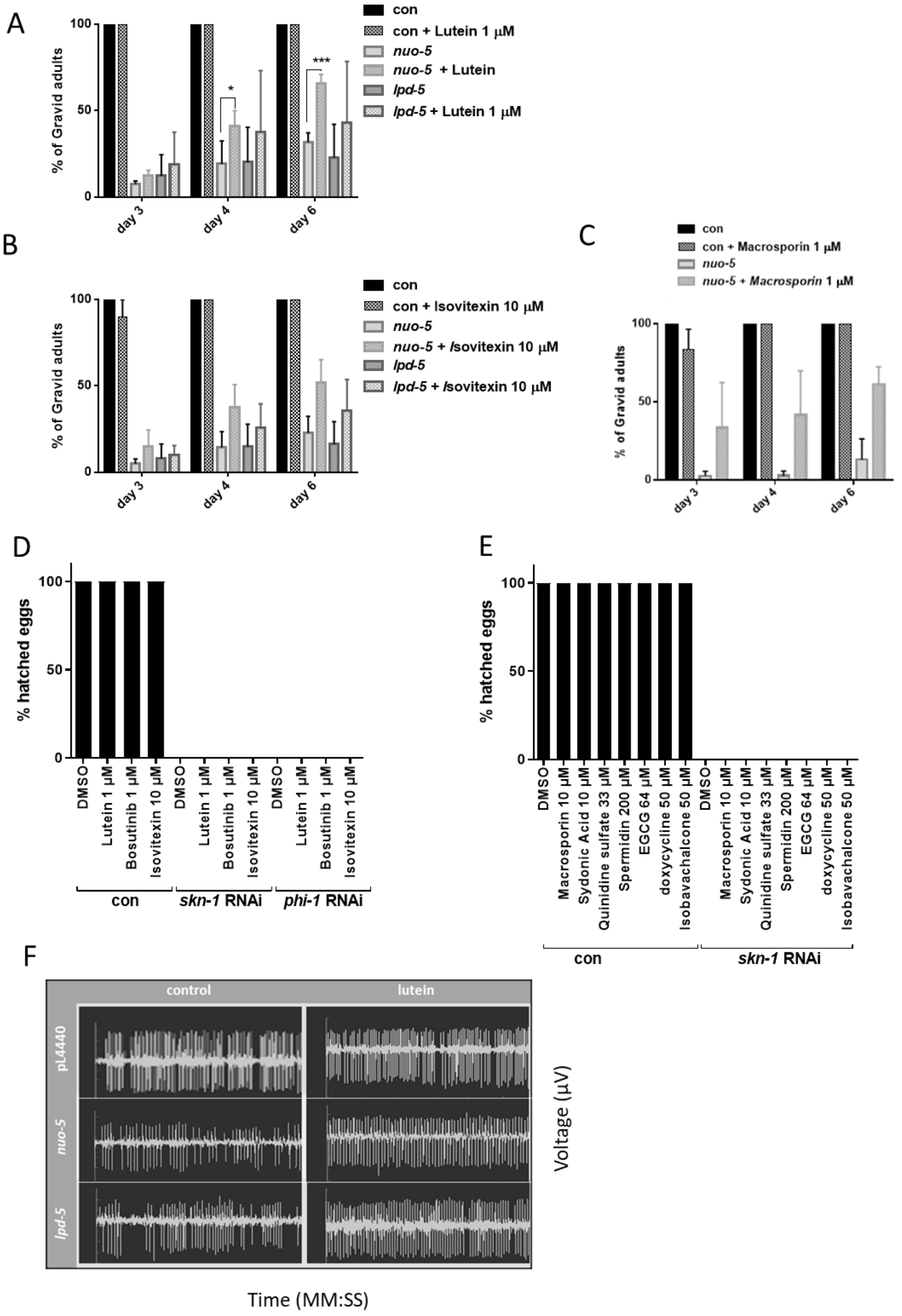

**Figure S7. Effects of selected drugs on developmental rate and pharyngeal pumping.**
**A-C)** Development of *nuo-5* and *lpd-5*-depleted worms treated with either 1µM lutein or DMSO (**A**), with 1µM isovitexin or DMSO **(B**), or with 1µM macrosporin or DMSO (**C**). **D-E)** The typical embryonic lethality induced *phi-7* (**D**-**E**) and *skn-1* (**E**) RNAi is conserved upon drugs treatment. **F**) Representative EPGs, 20 seconds frame of ~2 minutes recordings are shown for each condition. Pharyngeal pump frequency was measured using the ScreenChipTM System (InVivo Biosystems).

**Supplementary Figure 8**

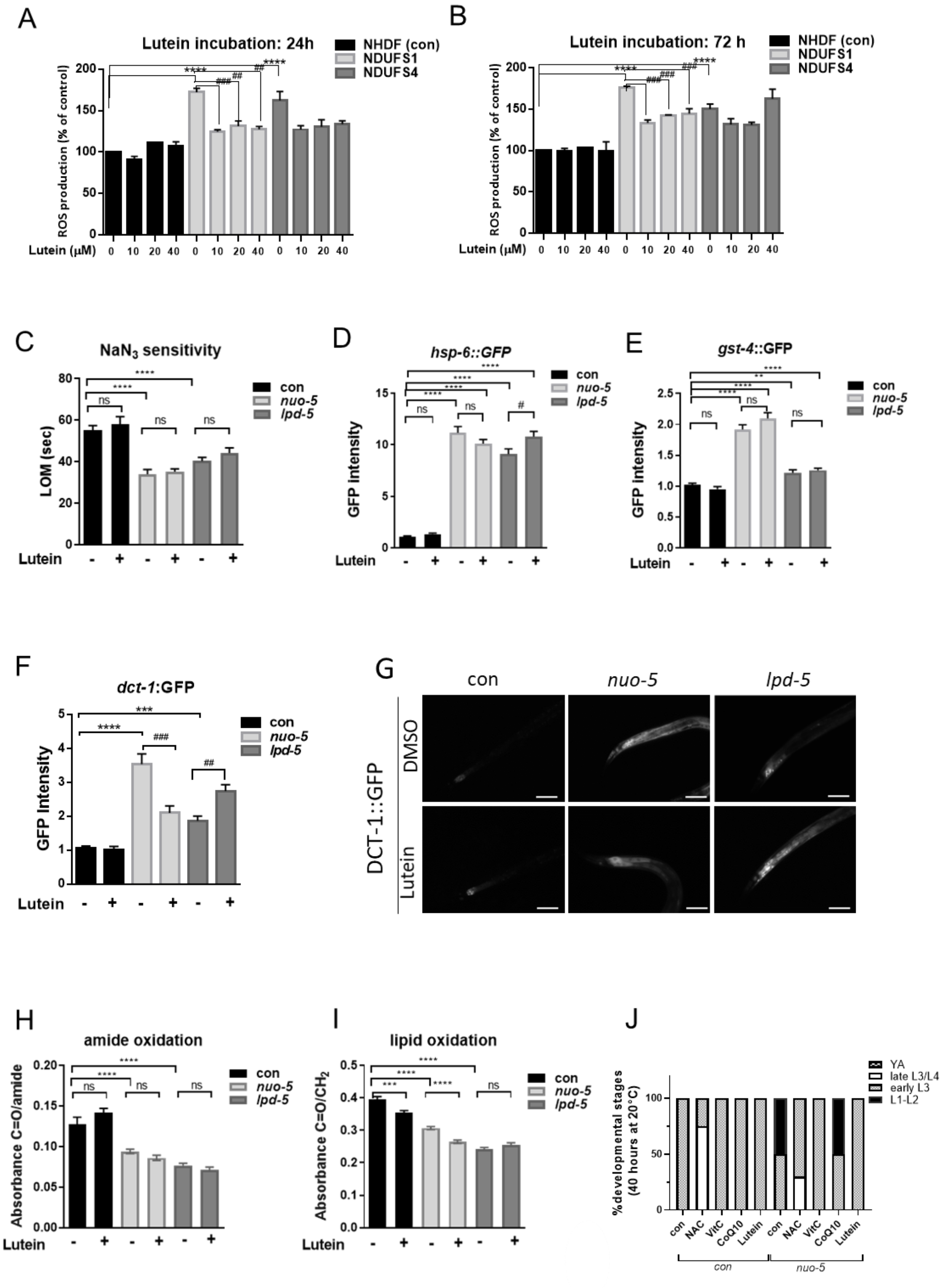


**Figure S8. Effects of lutein on different parameters in *nuo-5* and *lpd-5 C. elegans* mutants and patients' derived cells with mutations in the corresponding genes. A**) ROS production in human skin fibroblasts with lutein 24 hours incubation and **B**) 72 hours incubation. Different concentrations of lutein were used (N=3)**. C**) Sensitivity to 10 mM sodium azide, LOM = last observed movement. **D**) Quantification of GFP expression in WT strains expressing the $p_{hsp\text{-}6}$*::GFP*. **E)** Quantification of GFP expression in WT strains expressing the $p_{gst\text{-}4}$*::GFP*. **F**) Quantification of GFP expression in WT strains expressing the $p_{dct\text{-}1}$::DCT-1::GFP transgene. N=2, n=15-25 per group and corresponding representative pictures (**G**) Scale bar 50 µm**.** Animals are fed as in Figure 2 with and without lutein (1 µM final concentration) administration. **H**) Amide oxidation and **I**) lipid oxidation, both assessed with SR-µFTIR. N=3, n=15-20 per group. **J)** Comparison of development between control and worms fed nuo-5 RNAi treated either with DMSO or the indicated antioxidant. Bars indicate the percentage of different developmental stages after 40 hours from egg lay. L1 -L2, 1st to 2nd larval stage; early L3, beginning of 3rd larval stage; late L3/L4, end of 3rd larval stage and beginning of 4th larval stage. N=2, n=40. One-way ANOVA, asterisks (*) denote significant differences vs control, # denote differences among conditions. *p-value <0.05 ** p value < 0.01, ***p-value < 0.001 ****p-value < 0.0001. Bar graphs represent means ± SEM.

**Supplementary Figure 9**

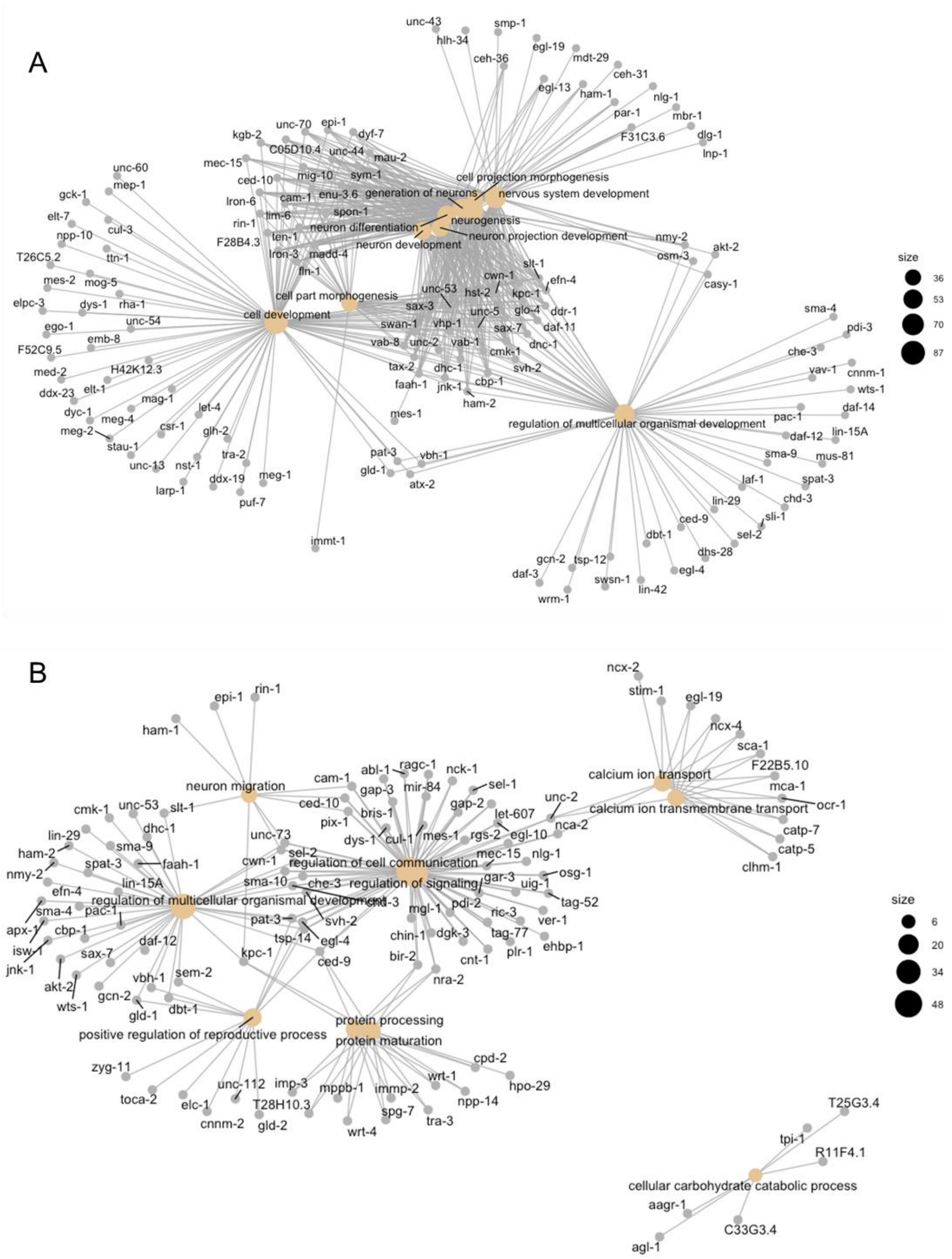


**Figure S9. Biological complexity network. A)** Microarray analysis comparing control vs *nuo-5* strong treated animals (results showed in **Fig 5**); **B**) control vs *nuo-5* strong + lutein. Created in R, using Bioconductor.

**Supplementary Figure 10**

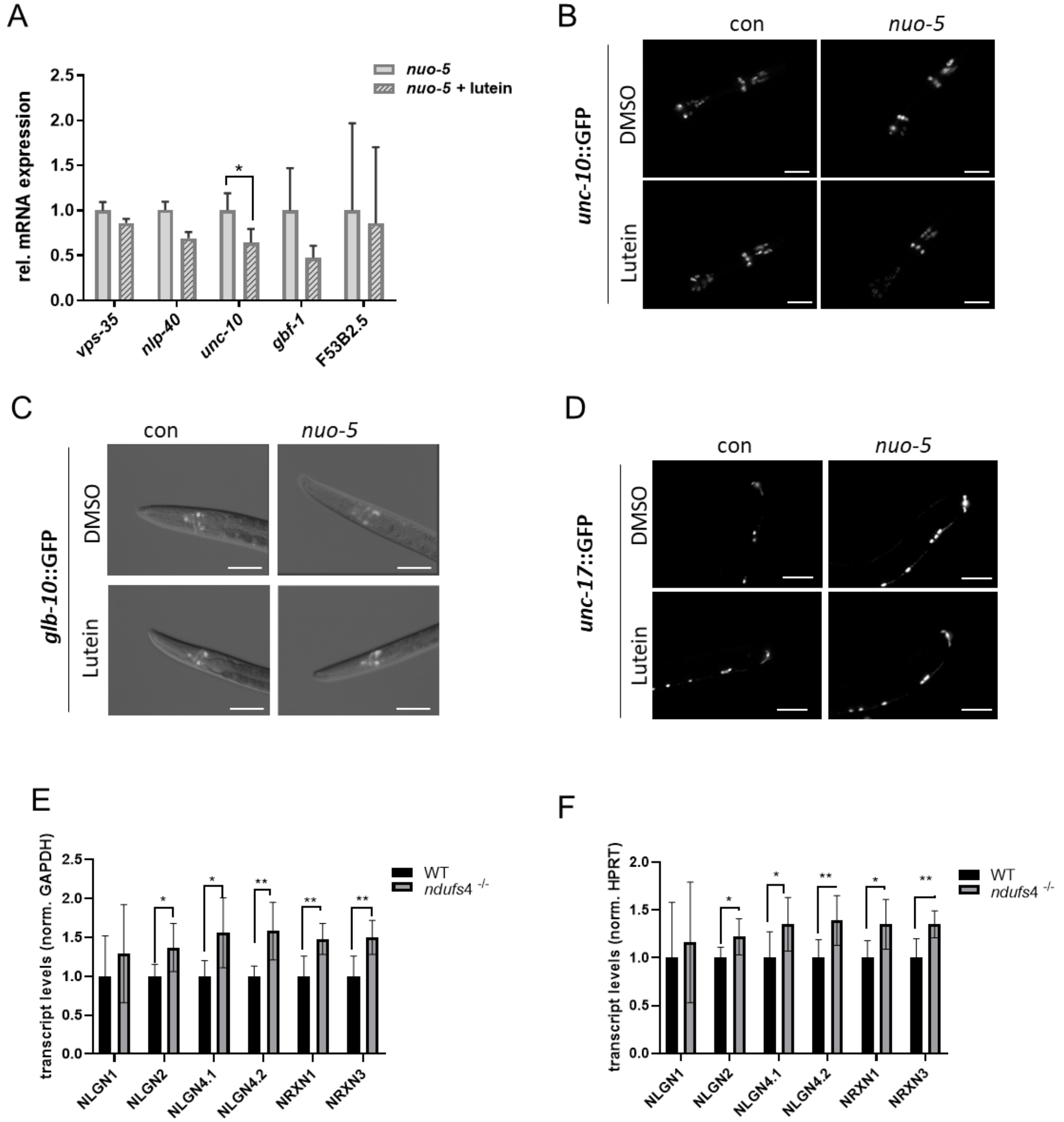


**Figure S10. Lutein affects synaptic genes' expression.**

**A)** Relative mRNA expression of *vps-35*, *nlp-40*, *unc-10*, *gbf-1* and F53B2.5 assessed by qPCR. **B**) Representative pictures of $p_{unc\text{-}10:}$:GFP in head neurons. Scale bar 20 µm. **C)** Representative pictures of $p_{glb\text{-}10}$::GFP in head neurons. Scale bar 50 µm. **D**) Representative pictures of $p_{unc\text{-}17}$::GFP in head neurons. Scale bar 50 µm. **E-F)** Expression levels of neuroligin (NLGN1-2 and 4) and neurexin (NRX1 and 3) transcripts in brains of NDUFS4-/- mice compared to healthy animals, normalized to **E**) GAPDH or **F**) HPRT**,** assessed by qPCR**.** One-way ANOVA asterisks (*) denote significant differences vs control, # denote differences among conditions. *p-value <0.05 ** p value < 0.01, ***p-value < 0.001, ****p-value < 0.0001. Bar graphs represent means ± SEM.

#### Supplementary Figure 11

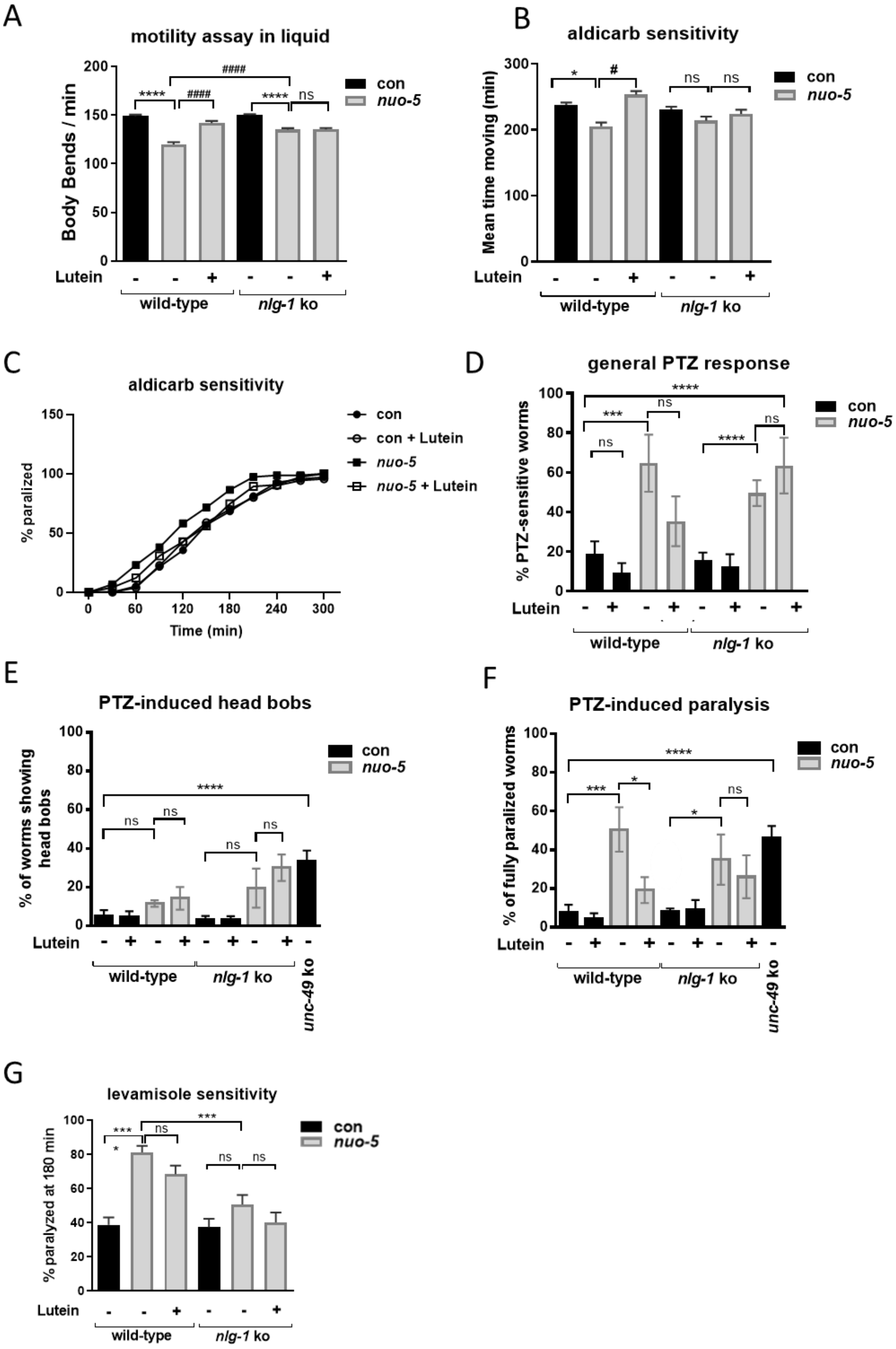

**Figure S11. Lutein effects are mediated by *nlg-1* suppression.** **A**) Frequency of lateral swimming ("thrashing") movements quantified manually in liquid media for one minute. N=3, n=15.

**B**) Aldicarb-induced paralysis curves and **C**) mean time moving on aldicarb plates of control and *nuo-5* in the presence or absence of lutein. (Aldicarb concentration: 0.5 mM) N=3, n=25-30 per group. **D)** PTZ general response, comprehensive of **E**) PTZ-induced head bobs. N=4, n=15-17. Bars indicate the percentage of animals showing head convulsions (head bobs) phenotypes after 1 hour of exposure to 2.5 mg/ml PTZ; and **F**) PTZ induced full body paralysis. N=4, n=15-17. Bars indicate the percentage of animals showing full body paralysis after 1 hour of exposure to 2.5 mg/ml PTZ. The *unc-49* knockout strain was used as positive control. **G**) Sensitivity to Levamisole. Bars indicate mean survival on plates containing 50 µM Levamisole. N=4, n=25-30. One-way ANOVA asterisks (*) denote significant differences vs control, # denote differences among conditions. *p-value <0.05 ** p value < 0.01, ***p-value < 0.001, ****p-value < 0.0001. Bar graphs represent means ± SEM.

**Supplementary Figure 12**

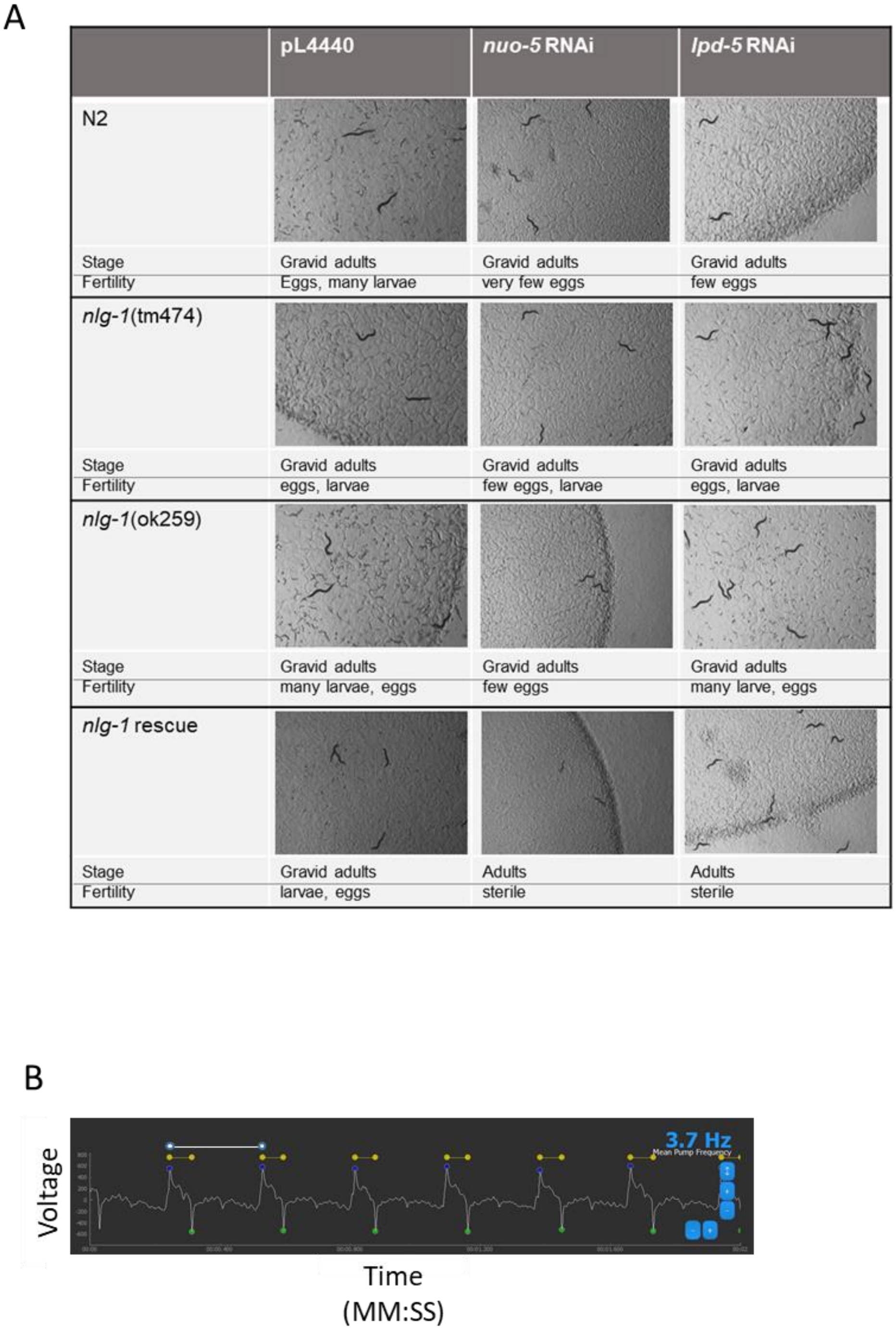


**Figure S12. Effects of gene silencing on development of RNAi hypersensitive strains and EPG representative track.**

**A)** Representative pictures and brief developmental description of different C. *elegans* strains treated with mild RNAi against *nuo-5* or *lpd-5* (parental generation), after 4 days from egg-lay.

*nlg-1* rescue corresponds to CRR104 transgenic strain (crrEx4 [pPD95.77 (*Pnlg-1*::*nlg-1*); pDD04NeoR (P*myo*-2::GFP)]. Pictures were acquired with a Leica MZ10 F modular stereo microscope connected to a color digital camera. Scale bars 500 µm.
**B)** Representative EPG recording of a 5-days old worm. The voltage change is plotted against the time. Upon contraction of the pharynx, a positive voltage change can be observed as an excitatory spike (E spike, shown as blue dots). The relaxation of the pharynx muscle results in an adverse voltage change, the relaxation spike (R spike, shown as green dots). The horizontal yellow line represents the pump duration while the white line shows an inter-pump interval (IPI), the interval between one E spike and the next neighboring E spike. The R/E ratio is the ratio of the repolarization and depolarization waves in each pump (R and E waves) and is a parameter that indicates probable alterations in the normal muscle contraction/relaxation.

#### Supplementary Tables

#### Supplementary Table 1. List of genes whose different degree of silencing did not lead to consistent change in the observed phenotypes

| Cosmid ID | Gene | Human ortholog, short gene description | Disease |
|---|---|---|---|
| F08A8.1 | F08A8.1 | ACOX-1, Acyl-CoA oxidase, complex IV | Pseudo-natal adrenoleukodystrophy |
| B0432.2 | *djr-1.1* | PARK7 | Parkinson’s disease |
| T24H7.1 | *phb-2* | PHB2 (prohibitin 2). Is involved in several processes, including defecation; gamete generation; and mitochondrion morphogenesis | |
| EEED8.9 | *pink-1* | PINK1 (PTEN-induced kinase) | Parkinson’ disease |
| ZK1248.14 | *fzo-1* | Mfn-2, mitofusin1-GTPase/ | Charcot Marie Tooth (CMT2) |
| F41G3.4 | *fis-1* | fis-1, fission1, collaborate with DRP-1(dinamin related protein) in the mitochodniral fisison process | optic dystrophy |
| T05H10.6 | *pdha-1* | PDHA1 (pyruvate dehydrogenase E1 subunit alpha 1) and PDHA2 (pyruvate dehydrogenase E1 subunit alpha 2) | pyruvate decarboxylase deficiency |
| F42A8.5 | *sdhb-1* | SDHB, tag-55, Fe-S complex II | extradrenal pheocromocitoma |
| Y57A10A.15 | Y57A10A.15 | POLG, mtDNA pol | PEO-progressive exteranl ophtalmoplegia |
| T24C4.1 | *ucr-2.3* | UQCRC2 (ubiquinol-cytochrome c reductase core protein 2). Predicted to have metalloendopeptidase activity | mitochondrial complex III deficiency nuclear type 5 |
| R10E4.5 | *nth-1* | NTHL1 (nth like DNA glycosylase 1) | familial adenomatous polyposis 3 |
| M03C11.5 | *ymel-1* | YME1L1 (YME1 like 1 ATPase) | optic atrophy 11 |
| F45E4.9 | *hmg-5* | TFAM (transcription factor A, mitochondrial). Is involved in mitochondrial DNA metabolic process. | |
| T01B11.4 | *tag-316* | SLC25A4 (solute carrier family 25 member 4) and SLC25A5 (solute carrier family 25 member 5) | |
| Y43C5A.5 | *thk-1* | TK1 (thymidine kinase 1). Is predicted to have thymidine kinase activity | |
| Y38F2AR.7 | *ppgn-1* | SPG7 (SPG7 matrix AAA peptidase subunit, paraplegin) | hereditary spastic paraplegia 7 |
| Y22D7AL.5 | *hsp-6* | HSPA9 (heat shock protein family A (Hsp70) member 9) | autosomal dominant sideroblastic anemia 4 |

| | | | |
|---|---|---|---|
| F46E10.8 | *ubh-1* | UCHL1 (ubiquitin C-terminal hydrolase L1) and UCHL3 (ubiquitin C-terminal hydrolase L3) | |
| F13B9.8 | *fis-2* | FIS1 (fission, mitochondrial 1). Is involved in apoptotic process | |
| C03G5.1 | *sdha-1* | SDHA (succinate dehydrogenase complex flavoprotein subunit A) | dilated cardiomyopathy 1GG; mitochondrial complex II deficiency; and paraganglioma |
| F21G4.6 | F21G4.6 | HTT (huntingtin) | Huntington's disease |

**Supplementary Table 2. Lifespan summary statistics undiluted and diluted**

| Gene/ RNAi clone | RNAi power | Mean Lifespan ± SE (Days) | Lifespan change (%)[a] | P value vs control[b] | P value mild vs strong[c] | Sample size/ n trials |
|---|---|---|---|---|---|---|
| con | | 18,61 ± 0,26 | | | | 330/4 |
| *spg-7* | mild<br>strong | 22,79 ± 0,34<br>21,69 ± 0,32 | +22,4 [1]<br>+16,5 | < 0,0001<br>< 0,0001 | <br>0,023 | 230/3<br>230/3 |
| con | | 18,13 ± 0,27 | | | | 250/3 |
| *nuo-1* | mild<br>strong | 23,91 ± 0,51<br>27,83 ± 0,68 | +31,8<br>+53,5 | < 0,0001<br>< 0,0001 | <br>< 0,0001 | 200/3<br>200/3 |
| T20H4.5 | mild<br>strong | 18,74 ± 0,32<br>27,34 ± 0,66 | +3,36<br>+50,79 | 0,63<br>< 0,0001 | <br>< 0,0001 | 160/2<br>140/2 |
| *dlst-1* | mild<br>strong | 18,49 ± 0,29<br>20,37 ± 048 | +1,98<br>+12,34 | 2,13<br>< 0,0001 | <br>0,001 | 200/3<br>200/3 |
| con | mild | 17,83 ± 0,56 | | | | 250/3 |
| *pdr-1* | mild (1:10)<br>mild (1:50)<br>strong | 17,75 ± 0,27<br>20,56 ± 0,47<br>13,38 ± 0,22 | - 0,44<br>+15,31 [1]<br>-24,95 | 5,32<br>< 0,0001<br>< 0,0001 | <br>< 0,0001<br>< 0,0001 | 200/3<br>80/1<br>200/3 |
| *dlat-1* | mild<br>strong | 18,41± 0,26<br>17,59 ± 0,30 | +3,25 [1]<br>-1,34 | 0,2236<br>0,76 | <br>0,19 | 200/3<br>200/3 |
| con | | 17,70 ± 0,28 | | | | 235/3 |
| *tag-61* | mild<br>strong | 21,54 ± 0,36<br>22,40± 0,41 | +21,46 [1]<br>+ 26,55 | < 0,0001<br>< 0,0001 | <br>0,73 | 250/3<br>220/3 |
| con | | 17,55 ± 0,34 | | | | 190/3 |
| *hmgs-1* | mild (1:10)<br>mild (1:50)<br>strong | 11,32 ± 0,45<br>14,54 ± 0,95<br>8,07 ± 0,25 | -35,49 [1]<br>-17,15<br>-54,0 | < 0,0001<br>< 0,0001<br>< 0,0001 | <br>< 0,0001<br>< 0,0001 | 140/2<br>80/1<br>140/2 |

All these clones showed a strong phenotype already in the parental generation (P0, see Figure 1) when used undiluted, and the mild effect was thus achieved by diluting the dsRNA expressing bacteria (P0, 1/10 or 1/50).

[a] % increase normalized mean lifespan compared to control; [b] Kaplan-Meier survival analysis, Log-rank test against control; [c] Kaplan-Meier survival analysis, Log-rank test between mild and strong treatment.

**Supplementary Table 3. Lifespan summary statistics P0 and F1.**

| Gene/ RNAi clone | RNAi power | Mean Lifespan ± SE (Days) | Lifespan change (%)[a] | P value vs control[b] | P value mild vs strong[c] | Sample size/ n trials |
|---|---|---|---|---|---|---|
| con | mild<br>strong | 19,05 ± 0,28<br>19,98 ± 0,40 | | | 0,61 | 275/4<br>140/3 |
| *lpd-5* | mild<br>strong | 22,01 ± 0,36<br>23,44 ± 0,65 | +15,53 [1]<br>+17,31 | < 0,0001<br>< 0,0001 | 0,0033 | 220/3<br>190/3 |
| *sco-1* | mild<br>strong | 20,04 ± 0,41<br>26,36 ± 0,8 | +4,38 [1]<br>+31,8 | 0,062<br>< 0,0001 | < 0,0001 | 220/3<br>200/3 |
| *nuo-5* | mild<br>strong | 24,91 ± 0,62<br>26,85 ± 1,00 | +30,76 [1]<br>+34,38 | < 0,0001<br>< 0,0001 | 0,125 | 200/3<br>200/3 |
| con | mild<br>strong | 20,3 ± 0,27<br>19,46 ± 0,38 | | | 0,9 | 204/3<br>220/3 |
| *spg-7** | mild<br>strong | 23,69 ± 0,39<br>25,23 ± 0,35 | +16,69 [1]<br>+29,6 | < 0,0001<br>< 0,0001 | 0,0001 | 230/3<br>220/3 |
| *aldo-2* | mild<br>strong | 19,82 ± 0,40<br>18,50 ± 0,66 | -5,25<br>-6,56 | 0,08<br>0,70 | 0,652 | 160/2<br>140/2 |
| con | mild<br>strong | 21,65 ± 0,42<br>21,31 ± 0,42 | | | 0,73 | 235/3<br>235/3 |
| *sod-1* | mild<br>strong | 19,72 ± 0,31<br>18,65 ± 0,31 | - 9,91 [1]<br>- 12,48 | < 0,0001<br>< 0,0001 | 0.0184 | 240/3<br>240/3 |
| con | mild<br>strong | 23,60 ± 0,56<br>23,59 ± 0,55 | | | 0.83 | 160/2<br>150/2 |
| *cox-10* | mild<br>strong | 21,92 ± 0,47<br>21,50 ± 0,46 | - 7,11 [1]<br>- 8,85 | 0,01<br>0,0044 | 0,69 | 160/2<br>160/2 |
| con | mild<br>strong | 19,99 ± 0,33<br>21,87 ± 0,37 | | | <0,0001 | 235/3<br>220/3 |
| T20H4.5* | mild<br>strong | 26,18± 0,58<br>29,28± 0,65 | +30,96 [1]<br>+ 33,88 | < 0,0001<br>< 0,0001 | <0,0001 | 250/3<br>220/3 |
| con | mild<br>strong | 18,74 ± 0,34<br>21,49 ± 0,63 | | | 0,012 | 200/3<br>120/2 |
| *eat-3** | mild<br>strong | 21,33 ± 0,45<br>23,77 ± 0,95 | + 13,82 [1]<br>+ 10,60 | < 0,0001<br>< 0,0001 | 0,0027 | 140/2<br>120/2 |
| *aco-2* | mild<br>strong | 16,88 ± 0,44<br>17,46 ± 1,03 | - 9,92 [1]<br>- 18,75 | 0,013<br>0,027 | 0,28 | 140/2<br>120/2 |

| | | | | | | |
|---|---|---|---|---|---|---|
| con | mild | 21,15 ± 0,42 | | | | 150/2 |
| | strong | 20,19 ± 0,41 | | | 0,05 | 160/2 |
| *fum-1* | mild | 14,56 ± 0,23 | - 31,15 [1] | < 0,0001 | | 160/2 |
| | strong | 18,62 ± 0,33 | - 7,77 | 0,0006 | < 0,0001 | 160/2 |
| con | mild | 18,93 ± 0,42 | | | | 150/2 |
| | strong | 18,38 ± 0,45 | | | 0,57 | 160/2 |
| *drp-1* | mild | 18,61 ± 0,40 | -1,69 [1] | 0,37 | | 140/2 |
| | strong | 18,15 ± 0,36 | -1,25 | 0,35 | 0,425 | 140/2 |
| con | mild | 18,17 ± 0,29 | | | | 260/3 |
| | strong | 18,76 ± 0,45 | | | 1,16 | 140/2 |
| *ogdh-1* | mild | 18,69 ± 0,38 | +2,86 [1] | 2,89 | | 200/3 |
| | strong | 21,45 ± 0,50 | +14,33 | 0,0003 | < 0,0001 | 200/3 |
| con | mild | 20,69 ± 0,24 | | | | 315/4 |
| | strong | 20,29 ± 0,31 | | | 0,68 | 300/4 |
| F53F4.10 | mild | 23,73 ± 0,48 | +14,69 [1] | < 0,0001 | | 320/4 |
| | strong | 28,92 ± 0,74 | +42,53 | < 0,0001 | < 0,0001 | 300/4 |

[a] % increase normalized mean lifespan compared to control; [b] Kaplan-Meier survival analysis, Log-rank test against control; [c] Kaplan-Meier survival analysis, Log-rank test between mild and strong treatment. * These clones showed already a strong phenotype in the parental generation (P0, see Figure 1) when used undiluted, and a milder effect was thus also achieved by diluting the dsRNA expressing bacteria (See Table S2).

**Supplementary Table 4.**
**Summary of chemotaxis assays in *C. elegans* models of Complex I deficiency**

| RNAi clone | RNAi power | $NH_4Ac$ 2,5M (ASE) | Pyrazine (AWA) | Benzaldehyd (AWC) | Butanol (AWC) |
|---|---|---|---|---|---|
| ***nuo-2*** | mild | | ↑ | ↑ | |
| | strong | ↓ | ↓ | ↓ | ↓ |
| ***nuo-5*** | mild | | ↑ | | |
| | strong | ↓ | ↓ | ↓ | ↓ |
| **F53F4.10** | mild | | ↑ | | |
| | strong | ↓ | ≈ | ↓ | ↓ |
| ***nuo-1*** | mild | | | | |
| | strong | ↓ | | | |
| ***lpd-5*** | mild | | ↑ | ↑ | |
| | strong | ↓ | ↓ | ≈ | ↓ |
| **T20H4.5** | mild | | | | |
| | strong | ↓ | ↓ | ↓ | |

↑ Increased Chemotaxis index; ↓ decreased Chemotaxis index or ≈ no changes in the Chemotaxis index, compared to N2 worms fed bacteria expressing the empty-vector (pL4440)
Stage of the worms: L3 Larvae

**Supplementary Table 5. Developmental rescue on *cep-1(lg12501)* mutants**

| *C. elegans* gene | Human gene | Rescuing strong phenotype with *cep-1*(lg12501) | Rescuing strong phenotype returning on Pl4440 |
|---|---|---|---|
| *lpd-5* | NDUFS4 | Partial rescue | Yes, returning on Pl4440 within 24 h |
| *nuo-5* | NDUFS1 | Partial rescue | Yes, returning on Pl4440 as eggs |
| F53F4.10 | NDUFV2 | ND | Yes, returning on Pl4440 within 24 h |
| *nuo-1* | NDUFV1 | Not rescued | Yes, returning on Pl4440 as eggs |
| T20H4.5 | NDUFS8 | Partial rescue | Yes, returning on Pl4440 as eggs |
| *nuo-2* | NDUFS3 | Partial rescue | Yes, returning on Pl4440 within 24 h |

ND not determined

**Supplementary Table 6. List of compounds used in the phenotype-based screening to identify Complex I-associated disease suppressors**

| Treatment | Activity/ tissue of action | Dose (µM) | Lifespan extension in *C. elegans* (dose) | Screening Results: % of gravid adult at day 6 in *nuo-5* (dose) * [SEM, n] | Screening Results % of gravid adult at day 6 in *lpd-5* (dose) * [SEM, n] |
|---|---|---|---|---|---|
| Dimethyl sulfoxide (DMSO) | solvent | 0.25% | no | 10.3 [5.25, 8] | 6.7 [1.8, 8] |
| Flavonoids and polyphenols | | | | | |
| 4,4'-dimethoxychalcone | Autophagy inducer [2] | 50 | yes (41,6 µM)[2] | 0 [ND,1] | 0 [ND,1] |
| 4- 4-methoxychalcone | ND | 50 | no (our observation) | 0 [ND,1] | ND |
| 4'-methoxychalcone | ND | 100, 1000 | no (our observation) | 0 [ND,1] | ND |
| EGCG | Antioxidant, anti-inflammatory [3] | 64, 640 | yes (200 µM) [3] | **22** (64 µM) [22.19, 3] 0 (640 µM) [ND,1] | 17.3 (64 µM) [7.5, 5], 0 (640 µM) [0, 2] |

| | | | | | |
|---|---|---|---|---|---|
| Hesperetin | Antioxidant, anti-inflammatory [4] | 200 | ND | 11.4 [10.5,1] | 2,9 [2.9,2] |
| Quercetin | Antioxidant, anti-inflammatory [5] | 100 | yes (100 µM) [6] | 6.7 [6.6,3] | ND |
| Resveratrol | autophagy inducer [7] | 100, 150 | yes (100 µM) [8] | **25.9** (100 µM) [13.8, 3]<br>0 (150 µM) [ND,1] | ND |
| <u>Other Natural Products</u> | | | | | |
| Helenalin | Anti-inflammatory [9] | 10,100 | ND | 1,4 (100 µM) [0.5, 2] | 0 (10 µM) [ND,1],<br>15,6 (100 µM) [27.13, 2] |
| Hexaprenylhydro-chinone | H,K-ATPase inhibitor [10] | 100 | ND | 0 [ND,1] | 0 [ND,1] |
| Homosekikaic Acid | Antioxidant [11] | 100 | ND | 0 [ND,1] | 0 [ND,1] |
| 2-Hydroxy-4-methoxyphenylaceto-nitrile | ND | 100 | ND | 4.2 [ND, 1] | 4.1 [ND, 1] |
| Hydroxysydonic Acid | Antibacterial [12] | 10, 100 | ND | 0 [ND,1] | **23.5** (10 µM) [11.8, 3]<br>0 (100 µM) [ND,1] |
| Hyperoside | Antibacterial and antifungal [13] | 100 | ND | 1.9 [1.1, 2] | 0.4 [ND, 1] |
| Indol-3-carboxylic acid | Antibacterial and anthelmintic [14] | 100 | ND | 0 [ND,1] | 18.3 [9.4, 3] |
| Isobavachalcone | Antibacterial, antifungal, anticancer, antioxidant [15] | 25, 50 | ND | 0 (25 µM) [ND,1]<br>0 (1.5 µM) [ND,1] | 0 (25 µM) [ND,1]<br>0 (1.5 µM) [ND,1] |
| Isovitexin | Antioxidant, anti-cancer, anti-inflammatory [16] | 10, 100 | ND | **39.6** (10 µM) [14.8, 3] | **39.6** (10 µM) [26.6, 3]<br>10 (53.3 µM) [ND, 1] |
| Kämpferol-3-rutinoside | Antioxidant [17] | 100 | ND | 0 [ND,1] | 1.7 [ND, 1] |
| Kaempferitrin | Antioxidant [18] | 100 | ND | 0 [ND,1] | 0.8 [ND, 1] |
| Kahalalide F | Antitumor [19] | 0.5, 1, 1.5 | yes (0.5 µM) (our observation) | 9.6 (0.5 µM) [4.9, 5]<br>3.3 (1 µM) [1.6, 2] | **29.2** (0.5 µM) [1.7,2]<br>3.3 (1 µM) [ND, 1]<br>12.7 (1.5 µM) [4] |

| | | | | | |
|---|---|---|---|---|---|
| **Kojic acid** | Antifungal [20] | 100 | ND | 1.7 [0.8, 2] | 0.8 [ND, 1] |
| **Kuanoniamine D** | Anticancer [21] | 1, 10 | ND | 0 [ND,1] | 11 [11, 3] |
| **Longamide B** | Trypanocidal and antileishmanial [22] | 10 | ND | 0 [ND,1] | 2.5 [ND, 1] |
| **Luffariellolid** | Anticancer [23] | 10 | ND | 0 [ND,1] | 0 [ND,1] |
| **Lupeol** | ND | 10 | ND | 0 [ND,1] | 1.6 [ND, 1] |
| **Lutein** | Antioxidant, anti-inflammatory [24] | 1, 10, 50 | yes (100 µM) (our observation) | **47.9** (1 µM) [19.4, 4]<br>**25** (10 µM) [ND, 1] | **51.5** (1 µM)<br>0 (10 µM) [ND,1]<br>**20.7** (50 µM) [5.1, 2] |
| **Macrosporin** | Antifungal [25] | 10 | ND | **26** [18.8, 2] | **28.7** [15.4, 3] |
| **Manzamine A** | ATPase [26] | 0.5, 5, 10 | ND | 0 [ND,1] | 0 [ND,1] |
| **Manzamine F1** | Anti-infective [27] | 100 | ND | 2 [ND, 1] | 11 [11, 3] |
| **8-OH-manzamien A*** | ND | 5,10 | ND | 0 (5 µM) [ND,1]<br>0 (10 µM) [ND,1] | 0 (5 µM) [ND,1]<br>0.8 (10 µM) [ND,1] |
| **Spermidine** | autophagy inducer [7] | 200, 300 | yes (200 µM) [28] | **31.4** (200 µM) [15.3, 3]<br>1.7 (300 µM) [ND, 1] | 0.8 (300 µM) [ND, 1] |
| Other compounds | | | | | |
| **Doxycycline** | antibiotic | 20, 30, 50 | yes (30µg/mL) [29] | 8.7 (30 M) [7.9, 2]<br>9.6 (20 µM) [7.1, 2]<br>**25** (50 µM) [15.5, 2] | 7.9 (20 µM) [5.4, 2]<br>0.8 (30 µM) [0.8, 3] |
| **PP2** | SRC kinase family inhibitors [30] | 50 | ND | 0 (50 µM) [ND,1] | ND |
| **Bosutinib** | SRC kinase family inhibitors [30] | 1, 10 | ND | 16.6 (1 µM) [ND, 1]<br>16.6 (10 µM) [ND, 1] | 3.3 (10 µM) [ND, 1]<br>16.6 (1 µM) [ND, 1] |
| **DBE-99** | nervous system [31] | 1, 10 | ND | 0 (1 µM) [ND,1]<br>6.6 (10 µM) [ND, 1] | 0 (1 µM) [ND,1]<br>0 (10 µM) [ND,1] |
| **Quinidine Sulfate** | nervous system [32] | 33, 55 | yes (33 µM) [32] | 1.6 (33 µM) [1.6, 2]<br>1.2 (55 µM) [0.4, 2] | **31.4** (33 µM) [18.9, 3]<br>1.2 (55 µM) [1.2, 2] |
| **LYC-30904** | ATPase modulator [33, 34] | 10 | yes (10 µM) [35] | 12.5 [6.25, 2] | 1.2 [1.25, 2] |

* In control condition (worms fed pL4440 expressing bacteria) adult animals at day 6 are 100%, in worms fed *nuo-5* expressing bacteria adults at day 6 are 11.9%, and worms fed *lpd-5* expressing bacteria adults at day 6 are 7.3%.
ND not determined
[SEM, n] indicate standard error mean and number of replicates for each compound.
In **bold** results ≥20% of gravid adults at day 6.

**Supplementary Table 7** is a separate Excel file**. Gene Ontology (GO) analysis of the microarray experiment shown in Fig5.**

**Supplementary Table 8** is a separate Excel file. **List of transcripts (columns) in the Unsupervised hierarchical clustering map shown in Fig5B**.

**Supplementary Table 9**
**Summary of results with lutein on *nuo-5* depleted animals**

| **Phenotype** | | **con** | ***nuo-5*** | ***nuo-5* + lutein** |
|---|---|---|---|---|
| Development | | | | |
| Pharyngeal pumping | Pump Frequency | | | |
| | Pump Duration | | | |
| | IPI Duration | | | |
| | R to E ratio | | | |
| Movement | In agar | | | |
| | In liquid | | | |
| Sensory function | Attraction to Pyrazine | | | |
| | Attraction to Butanol | | | |
| Paralysis | Aldicarb | | | |
| | Levamisole | | | |
| | PTZ - Head bobs | | | |
| | PTZ- Full body | | | |
| ATP levels | | | | |
| mtDNA damage | | | | |
| nDNA damage | | | | |
| Lipid/protein oxidation | | | | |
| Resistance to oxidative stress | | | | |
| ROS in elegans | | | | |
| Gene's expression | *nuo-5* | | | |
| | *hsp-6* | | | |
| | *gst-4* | | | |
| | *hlh-30* | | | |
| | *dct-1* | | | |
| | *nlg-1* | | | |

Green = wild type phenotype
Purple = affected
Orange = partial rescue
White = not determined

**Supplementary Table 10: Primers' list used for the qPCRs.**

| List of primers | | |
|---|---|---|
| ***C. elegans* gene** | **Forward primer (5'-3')** | **Reverse primer (5'-3')** |
| ***vps-35*** | GGAATACGACCGACCAGGAG | TGCATCCATGGTTTTTCCCTTAT |
| ***nlp-40*** | GTTGCGGTTTTTGCGGCTC | ATCCAAGGAATGTGTCGGCT |
| ***unc-10*** | GGGGAGCAGAAGGGAAAAGA | CCAAGAGGACCATCAGCGAG |
| ***gbf-1*** | GGCTCGGAGACAATACCAACT | ATCGGCAACCTCATTCAGCA |
| **F53B2.5** | ATGCCATCACCAGTAGCTCG | AAGCATGTCGATCCGTGTGG |
| ***nuo-5*** | TCGAGCCGTTTCTGAGGTTT | GTGGGGAGCGATCTCATTCA |
| ***eif-3C*** | ACCACTCTGCAAACCGTCAA | ACTCTTGAGTGTCATCCTTCAG |
| ***csq-1*** | GCCAGGATCAGTTGCTCTCA | AAGTTGGGGCGAAAAGTGGA |
| **Mouse gene** | | |
| **mNLGN1** | GGTACTTGGCTTCTTGAGCAC | AAACACAGTGATTCGCAAGGG |
| **mNLGN2** | TGTCATGCTCAGCGCAGTAG | GGTTTCAAGCCTATGTGCAGAT |
| **mNLGN3** | CCCTGGGCTTCCTCAGTTTG | GGCAATGGTACTCTGGCACC |
| **mNLGN4-1** | CCCAACGAAGATTGCCTCTAT [36] | TCCATCTTCCGTGGGCACATACAC |
| **mNLGN4-2** | CCCAACGAAGATTGCCTCTAT | CGTCATTATCCGCTAAGTCCTC |
| **mNRXN1** | AACGGACTGATGCTTCACACA | GATATTGTCACCTGACGCAGATT |
| **mNRXN2** | CGGCTCACCTGACGTTAAACA | TCCTTGCCTTTTGTCGGCTG |
| **mNRXN3** | GCACTATCCTACAGGCAACAC | GGCCGGTTATATTTGAAGGGGA |

#### Supplementary Methods

##### *C. elegans* strains

N2: wild-type (Bristol), $p_{glb-10}$::GFP; SJ4100:zcIs13[$p_{hsp-6}$::GFP], CL2166 dvIs19 [(pAF15)$p_{gst-4}$::GFP::NLS] III, JIN1679: jinEx10 [$p_{hlh-30}$::HLH-30::GFP + *rol-6*(su1006)], LX929:vsIs48 [$p_{unc-17}$::GFP], PE255:feIs5 [$p_{sur-5}$::luciferase::GFP + *rol-6*(su1006)] X, IR1431: N2;Ex001[$p_{dct-1}$DCT-1::GFP], AU78: agIs219 [T24B8.5p::GFP::unc-54-3' UTR + ttx-3p::GFP::unc-54-3' UTR] III [37], KP3928 nuIs165 [punc-129::UNC-10::GFP], VC228: nlg-1(ok259) X, vjIs47[$p_{nlg-1::}$GFP]IV, *vjIs105*[$p_{nlg-1}$*::nlg-1-gfp*]III [38], TU3401: sid-1(pk3321); uls69[myo2p::mCherry + unc-119p::sid-1], CL6114: nre-1(hd20) lin-15b(hd126), CRR100: *nlg-1*(ok259) X; crrEx4 [pD95.77; pDD04Neo$^R$ ($P_{myo-2}$::GFP)] [39], CRR104: *nlg-1*(ok259) X; crrEx4 [pD95.77 ($P_{nlg}$-1::nlg-1); pDD04Neo$^R$ ($P_{myo-2}$::GFP)] [39] and CB407: unc-49(e407) III [40].

##### Lifespan assay

Survival analysis started from hatching and was carried out at 20°C. Animals were scored as dead or alive and transferred every day on fresh plates during the fertile period, and then every other day or every 3 days until death. Worms were considered dead when they stop pharyngeal pumping and responding to touch. Worms that died because of internal bagging, desiccation due to crawling on the edge of the plates, or gonad extrusion were scored as censored. These animals were included in lifespan analyses up to the point of censorship and were weighted by half in the statistical analysis. We calculated mean lifespan, standard deviation of the mean, and P value (Mantel-Cox regression analysis) from Kaplan-Meyer survival curves of pooled population of animals coming from at least two independent replicas. For statistical analysis we used the Online Application for Survival analysis OASIS 2 [41].

##### Chemotaxis assays

Sodium azide ($NaN_3$) was used to anesthetize worms. It was placed on buffered agar 180 degrees opposite on a 10 cm dish; the attractant (or repellent) was then placed on one NaN3 spot, and ethanol (neutral odor for the worms in which chemical was diluted) on the other spot; a population of 80-100 age-synchronized animals was placed in the center of the testing plate and the number of worms at attractant and control was counted every 15 minutes for four hours to calculate the Chemotaxis Index (CI). CI=(A-B)/(A+B+C), where A is the number of worms at attractant, B is the number of worms at control and C is the number of animals which didn't reach any of the two spots at the end of the two hours. For a population of 3 days old, wild-type animals, a good CI is around 0.8 for attractants and –0.8 for repellents after two hours (CI=0 means no attraction, while CI=1 or –1 represent maximum attraction or repulsion, but there are always some animals that, also for a wild-type strain, remain randomly dispersed in the assay plate or reach the control spot instead of the attractant) [42].

**Mitochondrial Respiration**

Synchronized wild-type (*N2)* worms were grown at 20°C for two consecutive generations on bacteria expressing dsRNA against the genes of interest: *nuo-5* and *lpd-5*. Mitochondrial respiration from these animals was compared with stage-matched animals left untreated or fed for only one generation (mild treatment). The Seahorse Bioscience Flux Analyzer was programmed with the appropriate protocol [43]. Worms were rinsed off the plate and washed and the number of animals estimated (by counting the number of worms in 20µl drops under a dissecting microscope). The 24-well Seahorse Cartridge was loaded with nearly 250 nematodes pipetted into each well. The final volume of each well was brought to 525µl with unbuffered EPA water. Two wells per assay were filled with EPA water without worms and used as blanks. 75 µl of each drug were loaded into the designated injection ports of the seahorse cartridge. The final solution concentrations after injection of each drug being the following: 20µM DCCD (1% DMSO), 25µM FCCP (2% DMSO), and 10mM sodium azide. The OCR was normalized to the actual number of worms in each well and the values obtained were again normalized on the actual protein content per worms measured with a BCA protein assay after sonication of 2000 nematodes for each condition. The average and the standard error of OCR per worm across the five replicates for each sample were calculated and a minimum of 10 time points of 5 wells was used.

**ATP levels**

Worms at the desired developmental stage (L2/L3 larvae) were transferred to 15mL conical tube, spun at 2200 RCF and the supernatant was removed. Nematodes were resuspended in k-medium to a concentration of 1.0±0.2 nematodes/µl. 50-100 animals were pipetted into each well of a white 96-well plate (4-5 wells per treatment), and brought to a final volume 100uL with k-medium. K-medium alone was used for blank measurements. A plate reader was used to read first the GFP fluorescence (Emissions filter: 502nm; excitation filter: 485nm) and then the luminescence after injection of a luminescence buffer. The ATP production was then calculated dividing the luminescence signal by the GFP intensity.

**Mitochondrial DNA copy number, mitochondrial DNA and nuclear DNA damage**

Synchronized wild-type (*N2)* worms were grown at 20°C for two consecutive generations on bacteria expressing dsRNA against the genes of interest: *nuo-5 and, lpd-5.* nDNA and mtDNA from these animals was compared with stage-matched animals fed bacteria expressing the empty vector pL4440. Six worms were picked and pooled in a single tube per biological replicate, and three biological replicates were taken per treatment in three experiments separated in time.

This assay defines the control samples as undamaged and determines a lesion frequency in experimental samples based on any decrease in amplification efficiency relative to the control samples [44]. One nuclear genome target (9,3 kb) and one mitochondrial genome target (10,9 kb) were amplified. The DNA damage assay is able to quantitatively measure the number of polymerase-stalling lesions based on the amount of amplification obtained from the QPCR [45-47]. To calculate DNA copy number we utilized a real-time PCR assay in which by using a standard the actual DNA content can be calculate [48, 49]. This method is particularly advantageous because the standard curve is run along with the samples and the actual number of copies can be calculated.

**Quantification of Gene Expression through fluorescent transgene reporters**

For the SJ4100 and CL2166 strains the GFP intensity of the total body was quantified (**Fig S4A**). Concerning the JIN1679 strain, for each animal the number of nuclei showing HLH-30 nuclear translocation (between the pharynx and the vulva) (**Fig S4B**) were counted with a macro specifically customized with ImageJ. With the KP3928 strain the intensity of the fluorescence was quantified in the pharynx, specifically, two circular area were designed with ImageJ corresponding to anterior and posterior pharyngeal bulbs and the GFP intensity was quantified in the two areas (**Fig S10A**). For $P_{glb\text{-}10}$::GFP the intensity of the neuronal nuclei located in the pharyngeal area was quantified (**Fig S10B**). For the $P_{dct\text{-}1}$::GFP a circular region in the middle body was selected to quantify the GFP signal (**Fig S8G**). For the selection of the area and the quantification of the signal the software ImageJ (http://imagej.nih.gov/ij/) was used. For the LX929 the intensity of the second proximal head neuron was quantified (**Fig S10D**). With the vjIs47, vjIs105 and CRR104 strains the intensity of the fluorescence was quantified in the ventral nerve cord (VNC), specifically, a rectangular area was designed in the posterior half of the nematode, between the vulva and the tale, and the GFP intensity was quantified in the selected areas (**Fig 7E and Fig 8F**).

**RNA extraction for Microarray**

Worms at the desired larval stage (L3) were washed off three big agar plates spotted with the RNAi bacteria under study (*nuo-5, lpd-5* and pL4440 for control). The plates were supplemented either with 1 µM Lutein or 0.25 % DMSO for control. Individual *C. elegans* were collected into nuclease-free water and frozen at −80 °C until needed. Frozen worm pellets were re-suspended in lysis buffer and the same volume of stainless-steel beads (6.0 mm RNase free) was added. A TissueLyser II (Qiagen) was used with 2 shakes for 2 minutes at 30 Hz. Total RNA was prepared from each sample using the RNeasy Mini Kit and QIAshredder according to the manufacturer's instructions (Qiagen-74104 and 79654). Total RNA from 5 different biological replicas was extracted, on column purified to remove small RNA fractions, and treated with DNAse (RNeasy kit, Qiagen).

**Supplementary information references**

31. *TOXICOLOGICAL REVIEW OF 2,2',4,4',5-PENTABROMODIPHENYL ETHER (BDE-99)*. 2008.
32. Ye, X., et al., *A pharmacological network for lifespan extension in Caenorhabditis elegans.* Aging Cell, 2014. **13**(2): p. 206-15.
33. Johnson, K.M., et al., *Identification and validation of the mitochondrial F1F0-ATPase as the molecular target of the immunomodulatory benzodiazepine Bz-423.* Chem Biol, 2005. **12**(4): p. 485-96.
34. Cleary, J., et al., *Inhibition of the mitochondrial F1F0-ATPase by ligands of the peripheral benzodiazepine receptor.* Bioorg Med Chem Lett, 2007. **17**(6): p. 1667-70.
35. Maglioni, S., et al., *An automated phenotype-based microscopy screen to identify pro-longevity interventions acting through mitochondria in C. elegans.* Biochim Biophys Acta, 2015.
36. Zhang, B., et al., *Autism-associated neuroligin-4 mutation selectively impairs glycinergic synaptic transmission in mouse brainstem synapses.* J Exp Med, 2018. **215**(6): p. 1543-1553.
37. Shivers, R.P., et al., *Phosphorylation of the conserved transcription factor ATF-7 by PMK-1 p38 MAPK regulates innate immunity in Caenorhabditis elegans.* PLoS Genet, 2010. **6**(4): p. e1000892.
38. Staab, T.A., et al., *Regulation of synaptic nlg-1/neuroligin abundance by the skn-1/Nrf stress response pathway protects against oxidative stress.* PLoS Genet, 2014. **10**(1): p. e1004100.
39. Calahorro, F. and M. Ruiz-Rubio, *Functional phenotypic rescue of Caenorhabditis elegans neuroligin-deficient mutants by the human and rat NLGN1 genes.* PLoS One, 2012. **7**(6): p. e39277.
40. Thapliyal, S. and K. Babu, *Pentylenetetrazole (PTZ)-induced Convulsion Assay to Determine GABAergic Defects in Caenorhabditis elegans.* Bio Protoc, 2018. **8**(17).
41. Han, S.K., et al., *OASIS 2: online application for survival analysis 2 with features for the analysis of maximal lifespan and healthspan in aging research.* Oncotarget, 2016.
42. Hart, A.C., *Behavior* in *WormBook, The C. elegans Research Community*. 2006.
43. Luz, A.L., et al., *Seahorse-based analysis of cellular respiration in Caenorhabditis elegans.* in preparation.
44. Meyer, J.N., *QPCR: a tool for analysis of mitochondrial and nuclear DNA damage in ecotoxicology.* Ecotoxicology, 2010. **19**(4): p. 804-11.
45. Furda, A.M., et al., *Analysis of DNA damage and repair in nuclear and mitochondrial DNA of animal cells using quantitative PCR.* Methods Mol Biol, 2012. **920**: p. 111-32.
46. Furda, A., et al., *Quantitative PCR-based measurement of nuclear and mitochondrial DNA damage and repair in mammalian cells.* Methods Mol Biol, 2014. **1105**: p. 419-37.
47. Hunter, S.E., et al., *The QPCR assay for analysis of mitochondrial DNA damage, repair, and relative copy number.* Methods, 2010. **51**(4): p. 444-51.
48. Rooney, J.P., et al., *PCR based determination of mitochondrial DNA copy number in multiple species.* Methods Mol Biol, 2015. **1241**: p. 23-38.
49. Venegas, V. and M.C. Halberg, *Measurement of mitochondrial DNA copy number.* Methods Mol Biol, 2012. **837**: p. 327-35.
